# Genetic and phenotypic heterogeneity in *PNPT1*, *MYO15A*, *PTPRQ* and *SLC12A2* variants detected among hearing impaired assortative mating families in Southern India

**DOI:** 10.1101/2021.04.06.438557

**Authors:** Paridhy Vanniya. S, Jayasankaran Chandru, Justin Margret Jeffrey, Tom Rabinowitz, Zippora Brownstein, Mathuravalli Krishnamoorthy, Karen B. Avraham, Le Cheng, Noam Shomron, C. R. Srikumari Srisailapathy

**Affiliations:** Department of Genetics, Dr. ALM PG Institute of Basic Medical Sciences, University of Madras, Chennai- 600 113, India; Department of Cell and Developmental Biology, Sackler Faculty of Medicine, Tel Aviv University, Tel Aviv- 6997801, Israel; Department of Human Molecular Genetics and Biochemistry, Sackler Faculty of Medicine and Sagol School of Neuroscience, Tel Aviv University, Tel Aviv- 6997801, Israel; BGI Genomics, Shenzhen- 518083, China

**Keywords:** *PNPT1*, *MYO15A*, *PTPRQ*, *SLC12A2*, assortative mating, South Indian hearing impaired

## Abstract

Exome analysis was used to resolve the etiology of hearing loss (HL) in four South Indian assortative mating families. Six variants, including three novel ones, were identified in four genes: *PNPT1* p.Ala46Gly and p.Asn540Ser, *MYO15A* p.Leu1485Pro and p.Tyr1891*, *PTPRQ* p.Gln1336*, and *SLC12A2* p.Pro988Ser. Compound heterozygous *PNPT1* variants were associated with prelingual profound sensorineural hearing loss (SNHL), vestibular dysfunction and unilateral progressive vision loss in one family. In the second family, *MYO15A* variants in the myosin motor domain, including a novel variant, were found to be associated with prelingual profound SNHL. A novel *PTPRQ* variant was associated with postlingual progressive sensorineural/mixed HL and vestibular dysfunction in the third family, with mastoid bone hypopneumatization observed in one family member. In the fourth family, the *SLC12A2* novel variant was found to segregate with severe-to-profound HL causing DFNA78, across three generations. Our results suggest a high level of allelic, genotypic and phenotypic heterogeneity of HL in these families. This study is the first to report the association of *PNPT1*, *PTPRQ* and *SLC12A2* variants with HL in the Indian population.

## Introduction

Non-syndromic hearing loss (HL) is known for its high heterogeneity. To date, more than 128 genes have been associated with this type of sensory impairment (https://hereditaryhearingloss.org/), but genetic diagnosis is complicated by the inherent diversity. Previous studies have implicated *GJB2* mutations as the most common genetic cause of HL, while *CDH23*, *SLC26A4*, *TMC1* and *MYO15A* comprise the second most common tier of genes (Brownstein et al. 2011; Duman and Tekin 2012; Ganapathy et al. 2014; Vanniya et al. 2018; Amritkumar et al. 2018; Chandru et al. 2020). Although a number of studies have investigated specific auditory gene frequencies (Yan et al. 2016; Subathra et al. 2016; Vanniya et al. 2018; Amritkumar et al. 2018; Chandru et al. 2020; Kalaimathi et al. 2020), the complexity of HL in Indian families remains underexplored.

Assortative mating implies non-random preferences in the choice of mates, which can either be positive (deaf marrying deaf – DXD) or negative assortative mating (deaf marrying normal – DXN). In our study, South Indian assortative mating hearing impaired (HI) families serve as a model system for exploring various dimensions of HL in the population, such as the influence of consanguinity and genotype-phenotype correlations in the population. They also provide a means to investigate gene-interactions on the backdrop of complementary (normal hearing offspring) and non-complementary unions (HI offspring).

In the present study, exome analysis was used to resolve the HL genotypes in four assortative mating South Indian HI families. The results provide an insight into the genotypic and phenotypic heterogeneity in South Indian assortative mating HI families.

## Materials and methods

### Subjects

In a cohort of 113 DFNB1-negative HI individuals from 83 assortative mating families, four HI probands were previously reported to have DFNB12 with homozygous or compound heterozygous *CDH23* variants (Vanniya et al. 2018). Five unrelated, DFNB12-negative probands, who were heterozygous for *CDH23* variants were selected for exome sequencing. For one family, *DXN*CHM46, exome sequencing was performed for the proband (II-3) and his six additional family members (including four HI and two normal hearing individuals). Altogether, exome sequencing was carried out in eleven individuals in the present study.

A total of 16 (HI and normal hearing) individuals from these four families were subjected to segregation analysis, and 50 normal hearing subjects with no history of HL were selected as age, sex and ethnicity matched controls for the study.

This study was approved by the Institutional Human Ethical Committee, and the subjects were included after signing a written informed consent, according to the rules of the Declaration of Helsinki.

### Molecular analysis

About 5ml of peripheral blood was collected and DNA was extracted by the phenol-chloroform-isoamyl alcohol (PCI) method. Exome sequencing was performed on the Illumina platform, by outsourcing to Macrogen Inc., South Korea, and BGI Genomics, China. For exome data analysis, the reads were aligned to the human reference genome GRCh37 (hg19), using BWA (Li and Durbin 2009) and variant calling was done with GATK (McKenna et al. 2010). Variants with >5% allele frequency were excluded from further analysis. ANNOVAR was used for novel variant annotation (Wang et al. 2010). Candidate genes were shortlisted by considering multiple parameters including gene functional annotation, the HL-associated gene list from the HHL homepage (https://hereditaryhearingloss.org/), and the available literature. Selected variants were ranked according to their status reported in the Deafness Variation Database (http://deafnessvariationdatabase.org/) and MAF in ExAC, 1000 Genomes, and gnomAD databases.

Sanger sequencing using exon-specific primers (Supplementary Table 1) designed with Primer3Plus (Untergasser et al. 2007) was carried out for variant confirmation, segregation analysis and variant screening in controls. As a cost-effective approach, the PCR-restriction fragment length polymorphism (RFLP) method was used to screen the specific variants in the controls, wherever possible (Supplementary Fig. 1).

### Clinical phenotyping

The clinical phenotypes associated with HL were profiled in the consenting HI probands and their family members. Audiometry was performed by an audiologist, to record hearing thresholds (250 to 8000 Hz). Vestibular assessment by videonystagmography (VNG) and video head impulse tests (vHIT) was carried out by authorized clinicians. Temporal lobe computed tomography (CT) and magnetic resonance imaging (MRI) were carried out to assess structural abnormalities.

### *In silico* analysis

Various *in silico* tools were used to study the effect of the variants identified. Evolutionary conservation of the mutation sites was studied using the phyloP (Pollard et al. 2010), phastCons (Siepel et al. 2005), and Consurf webservers (Ashkenazy et al. 2016). SIFT (Kumar et al. 2009), PolyPhen-2 (Adzhubei et al. 2015), PROVEAN (Choi et al. 2012), Mutation Taster (Schwarz et al. 2014), MutPred2 (Pejaver et al. 2017) and I-Mutant2.0 (Capriotti et al. 2005) were used for pathogenicity prediction of the variants.

In addition, structural modelling was used to predict the structural deviations caused by the variants. Homology modelling was done using Modeller v9.14 (Sali et al. 2016), and threading using I-Tasser (Zhang 2008) when homology modelling was not feasible. Energy minimization of the generated models was done using YASARA energy minimization server (Kreiger et al. 2009). The PDBsum (Laskowski 2001), ProSA (Weiderstein and Sippl 2007) and ProQ (Wallner et al. 2003) webservers were utilized for structure validation. Visualization of the model and superimpositions were performed using Swiss PDB viewer (Geux and Peitsch 1997). RMSD and TM-scores were obtained using the TM-align webserver (Zhang and Skolnick 2005).

## Results

In this study, exome sequencing resolved the genetic etiology of five unrelated probands from South Indian assortative mating families: *DXD*CHE32, *DXD*PTM56, *DXN*CHE29 and *DXN*CHM46. Six variants in four genes were found to be associated with HL in these families (Table 1). None of the variants were detected in the 50 normal hearing controls screened, and were not found among the gnomAD South Asian population (https://gnomad.broadinstitute.org/). The HL genotypes and the associated phenotypes observed in these four families are detailed below.

**Table 1.** Hearing loss-associated variants identified by exome sequencing in the four families

| Gene | Nucleotide change | Protein change | Protein domain | gnomAD# | ClinVar | SIFT | PolyPhen2 | PROVEAN | Mutation taster | MutPred2 | I-mutant 2.0 | phyloP | phastCons | Consurf | ACMG classification |
| --- | --- | --- | --- | --- | --- | --- | --- | --- | --- | --- | --- | --- | --- | --- | --- |
| *PNPT1* | c.137C>G | p.Ala46Gly | Polynucleotide phosphorylase | - | SCV001366117 | Tolerated | Probably damaging | Neutral | Disease causing | 0.253 | Decrease stability | 3.338 | 0.018 | 1 | PP1 |
|  | c.1619A>G | p.Asn540Ser | Polynucleotide phosphorylase | - | SCV001366118 | Damaging | Benign | Damaging | Disease causing | 0.526 | Decrease stability | 2.775 | 1 | 8 | PP1 |
| *MYO15A* | c.4454T>C | p.Leu1485Pro | Myosin motor | 0.000 | SCV001366121 | Damaging | Probably damaging | Damaging | Disease causing | 0.946 | Decrease stability | 5.147 | 1 | 9 | PM1 |
|  | c.5673C>G | p.Tyr1891* | Myosin motor | - | SCV001366122† | - | - | - | Disease causing | - | - | 2.493 | 1 | 6 | PVS1 |
| *PTPRQ* | c.4006C>T | p.Gln1336* | Fibronectin type III-14 | - | SCV001366119† | - | - | - | Disease causing | - | - | 0.473 | 0.997 | 3 | PVS1 |
| *SLC12A2* | c.2962C>T | p.Pro988Ser | Carboxy terminal domain | - | SCV001366123† | Tolerated | Probably damaging | Neutral | Disease causing | 0.486 | Decrease stability | 5.48 | 1 | 2 | PM1 |

(†Novel variant; #gnomAD South Asian population; MutPred2 scores above 0.5 signify pathogenicity; phyloP scores represent nucleotide conservation ranging from -14 and +6 (negative scores indicate fast-evolving sites, positive scores indicate conserved sites); phastCons scores signify nucleotide conservation scores ranging between 0 and 1 (values close to 1 denote conserved residues); Consurf scores represent amino acid conservation scores ranging from 1 to 9 (1 is variable, 9 is conserved); ACMG classification: PP1- Supporting evidence of pathogenicity, PM1- Moderate evidence of pathogenicity, located in critical and well-established functional domain, PVS1- Very strong evidence of pathogenicity- null/nonsense variant)

The female proband (II-2) in the *DXD*CHE32 family has compound heterozygous variants c.137C>G, p.Ala46Gly/c.1619A>G, p.Asn540Ser (Fig. 1B) in the *PNPT1* gene (MIM: 610316), associated with DFNB70 (MIM: 614934). Both these variants are located in the polynucleotide phosphorylase domain of PNPT1, which plays an essential role in mitochondrial RNA transport and is important for mitochondrial functioning (Von Ameln et al. 2012).

**Fig. 1.**
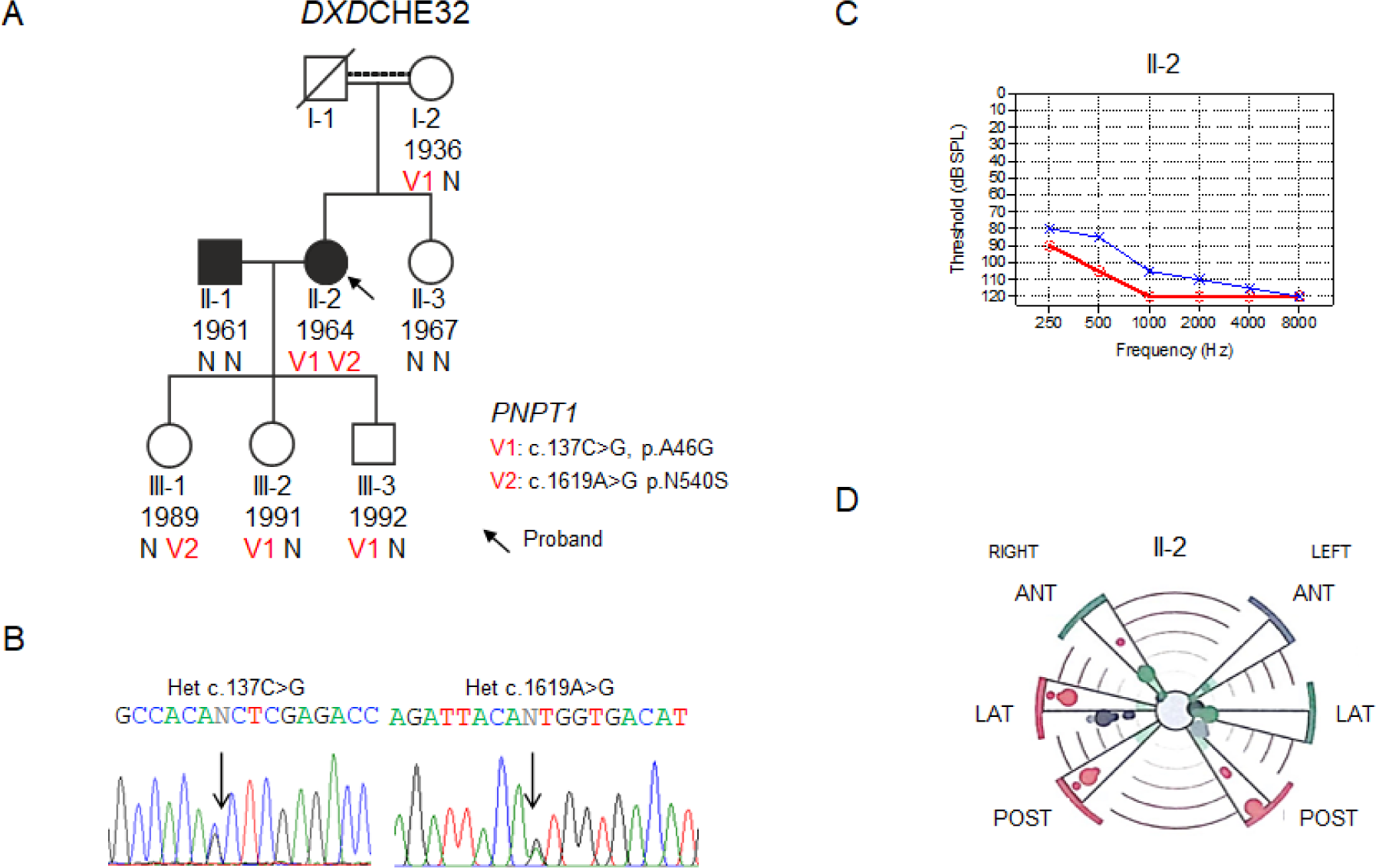
Segregation analysis and genotype-phenotype correlation in the *DXD*CHE32 family. The pedigree (A) shows the segregation of *PNPT1* variants, their chromatograms (B) and clinical phenotyping including audiogram (C) and vHIT (D) in the proband (II-2). Black filled symbols represent individuals with HL. V represents the variant allele and N the normal allele. The number under each individual is the birthdate. In some cases, the date is estimated due to lack of accurate records

The p.Ala46Gly variant is predicted to decrease protein stability (Table 1). The Ala46 residue is adjacent to the PNPT1 N-terminal mitochondrial transit peptide (spanning 45 amino acids); p.Ala46Gly may disturb the cleavage of this transit peptide. A comparison of the wild type (*wt*) and mutant (*mt*) protein structures suggests a predicted RMSD of 2.56 and TM-score of 0.92. p.Asn540Ser affects a highly conserved residue and is also predicted to reduce protein stability, with a RMSD of 2.55 and TM-score of 0.9 was predicted when comparing the *wt* and *mt* proteins.

Segregation analysis in the family revealed that the proband’s unrelated HI husband (II-1) is negative for both the *PNPT1* variants (complementary union), while their three normal hearing children are heterozygous for the two variants (Fig. 1A). The second normal hearing daughter (III-2) was found to be double heterozygous with *CDH23*:p.(Gly923Ser) and *PNPT1*:(p.Ala46Gly) (Table 2).

**Table 2.**
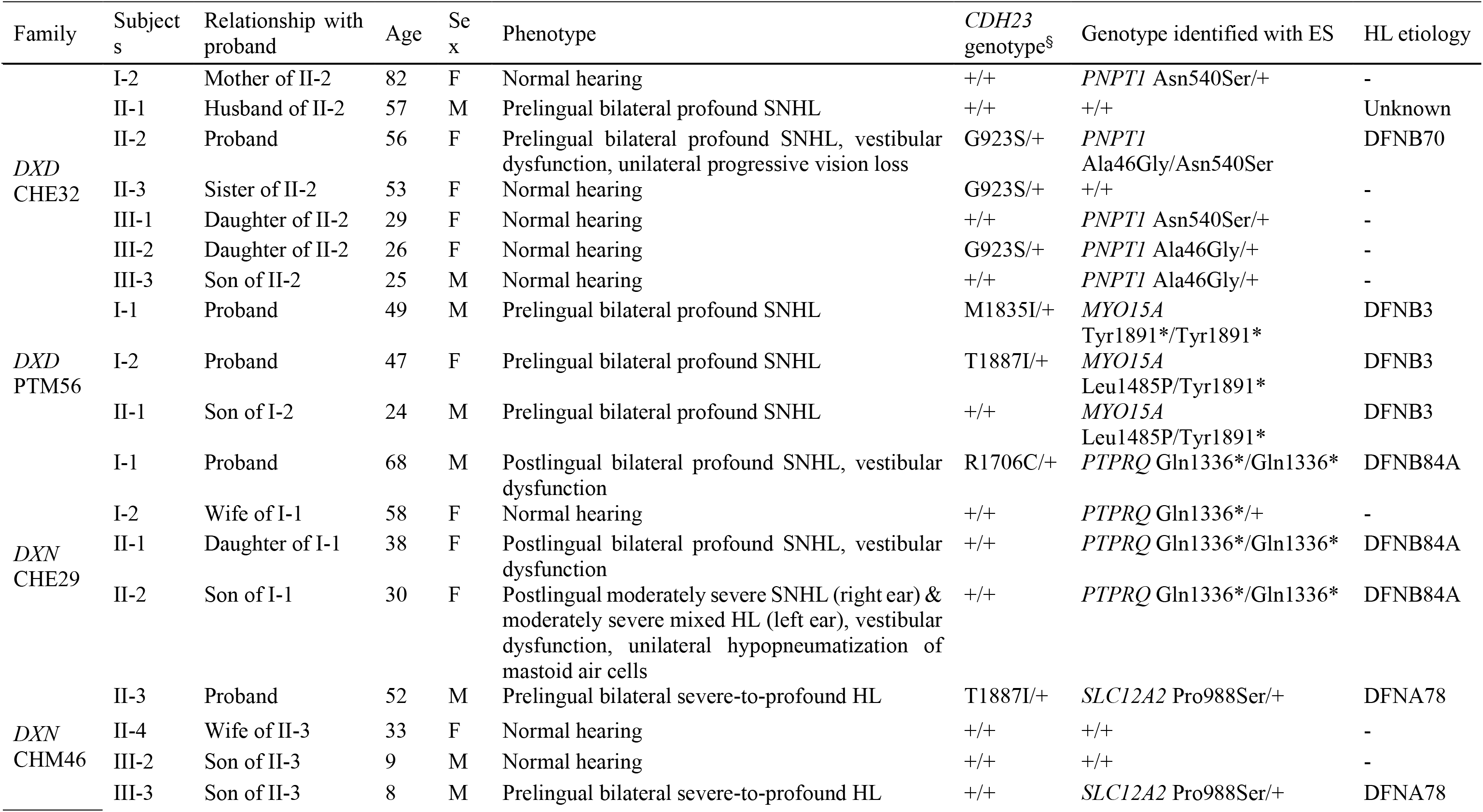
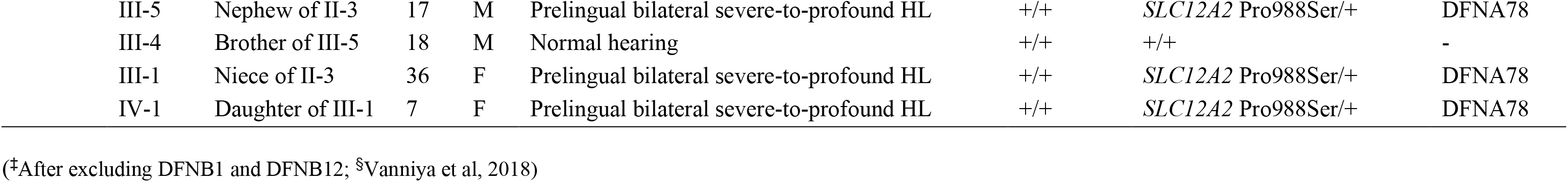
Hearing loss genotypes and associated phenotypes observed in the four families^‡^

The proband’s normal hearing parents are distantly related. Her mother (I-2) is heterozygous for *PNPT1* c.137C>G, p.Ala46Gly, and while her deceased father’s genotype could not be assessed, the proband’s normal hearing sister (II-3) tested negative for both variants (Table 2).

The proband exhibited prelingual bilateral profound sensorineural HL (SNHL) (Fig. 1C). VNG testing demonstrated gaze-evoked nystagmus with an inability to track optokinetic stimulations, suggesting a central lesion. vHIT revealed low saccades and vestibulo-ocular reflex (VOR) gain in the right anterior, lateral and posterior, and left posterior semi-circular canals (SCC), suggesting hypofunctional vestibules (Fig. 1D). The subject has progressive vision loss in the left eye and was diagnosed with diabetes 10 years ago.

Exome sequencing in family *DXD*PTM56 showed that both the unrelated husband (proband I-1) and wife (proband I-2) have *MYO15A* (MIM: 602666) variants associated with DFNB3 (MIM: 600316). While the husband is homozygous for a novel variant c.5673C>G, p.Tyr1891*, the wife (I-2) is compound heterozygous for the variants c.4454T>C, p.Leu1485Pro/c.5673C>G, p.Tyr1891* (Fig. 2B). Variants identified in both the probands occur in the myosin motor domain of Myosin XVa, which is encoded by *MYO15A*, and is required for actin organization at the stereocilia tips in the inner ear (Belyantseva et al. 2005).

**Fig. 2.**
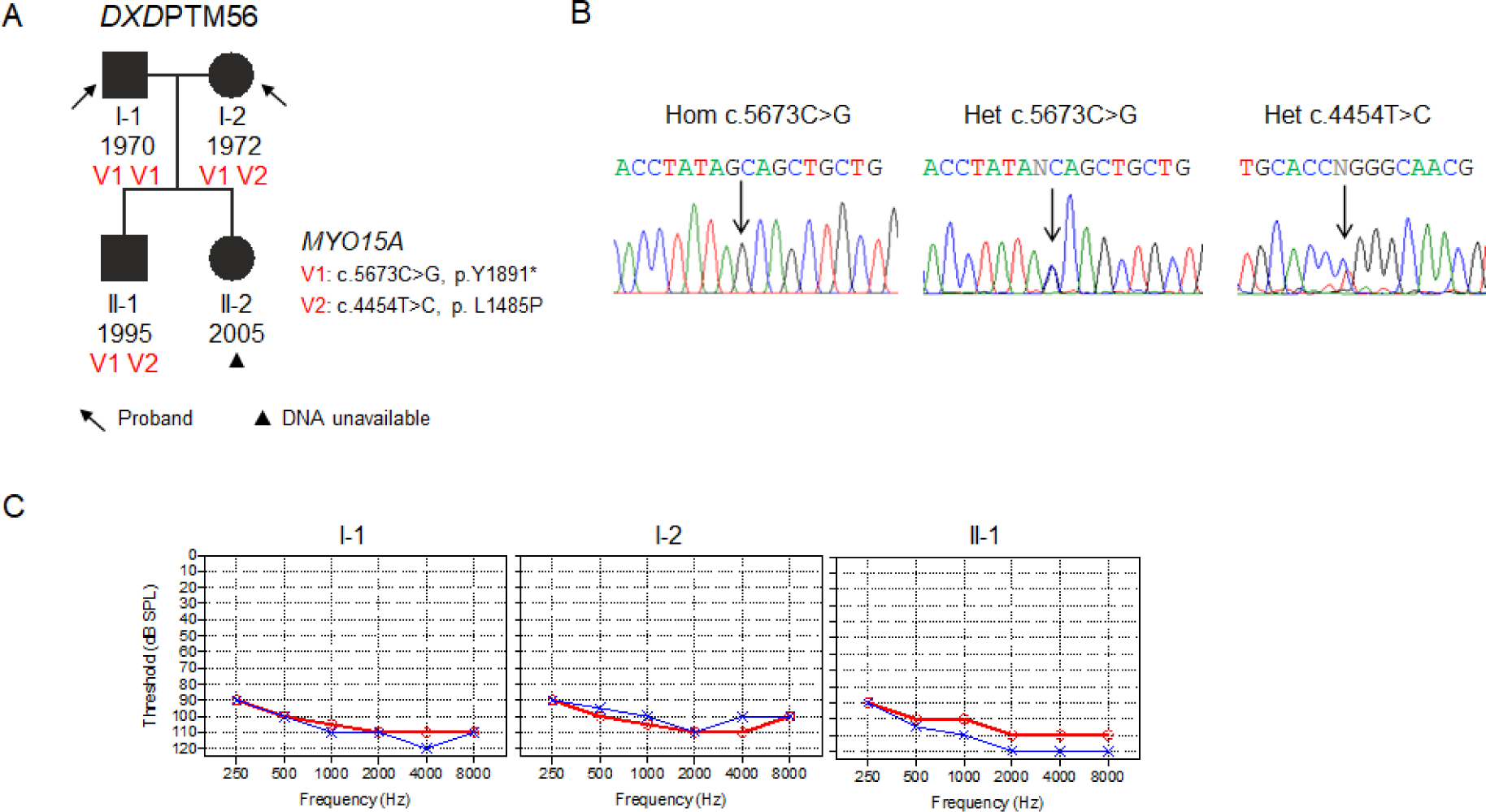
Segregation analysis and genotype-phenotype correlation in the *DXD*PTM56 family. The pedigree (A) depicts the segregation of *MYO15A* variants shown in the chromatograms (B), and (C) audiograms of I-1, I-2 (probands) and II-1

The p.Leu1485Pro variant is located in a highly conserved site and is predicted to reduce protein stability (Table 1). Structural analysis comparing *wt* and *mt* protein suggested an RMSD of 1.08 and TM-score of 0.96. p.Tyr1891* is predicted to lead to premature termination of the protein, resulting in the loss of the IQ-1, IQ-2, IQ-3, MyTH4-1, SH3, MyTH4-2 and FERM domains. An RMSD of 1.25 and TM-score of 0.9 were predicted when comparing the *wt* and *mt* proteins.

This being a non-complementary union, the couple’s two children (II-1 and II-2) are both HI. The HI son (II-1) was found to be compound heterozygous for *MYO15A* c.4454T>C, p.Leu1485Pro/c.5673C>G, p.Tyr1891* (Table 2), while the HI daughter (II-2) could not be tested due to non-consent (Fig. 2A). The HI probands (I-1 and I-2) and their son (II-1) had DFNB3, presenting with prelingual bilateral profound SNHL (Fig. 2C).

In the *DXN*CHE29 family, exome sequencing in the male proband (I-1) identified a novel homozygous variant c.4006C>T, p.Gln1336* (Fig. 3B) in *PTPRQ* (MIM: 603317), associated with DFNB84A (MIM: 613391). PTPRQ has phosphatase activity against phosphotyrosine and phosphatidylinositol molecules that are crucial for regulating cell survival (Seifert et al. 2003). This variant is located in the 14^th^ Fibronectin type-III (FN-III) domain of PTPRQ and is predicted to cause premature termination, leading to loss of four subsequent FN-III domains, a transmembrane domain and the tyrosine protein phosphatase domain. A comparison of *wt* and *mt* structural models predicts an RMSD of 2.63 and a TM-score of 0.85.

**Fig. 3.**
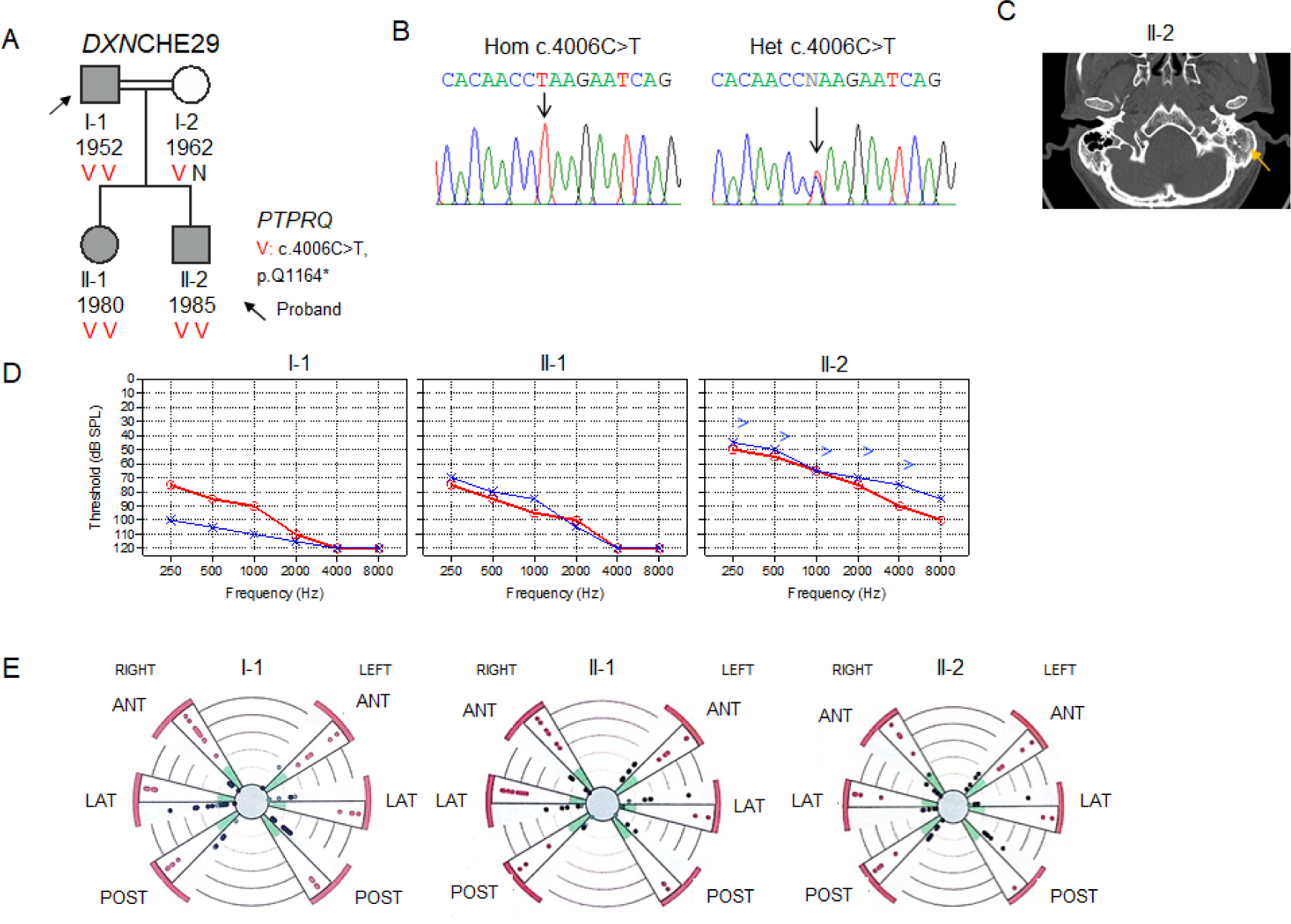
Segregation analysis and genotype-phenotype correlation in the *DXN*CHE29 family. The pedigree (A) shows segregation of postlingual progressive HL (grey filled symbols) with the *PTPRQ* variant seen in the chromatograms (B); the CT scan of II-2 indicating hypopneumatization in left mastoid air cells (C); the audiograms (D) and vHIT results of I-1, II-1 and II-2 (E)

The proband (I-1) in this family is a first cousin to his normal hearing wife (1-2) and also had parental consanguinity (Supplementary Fig. 2). Segregation analysis revealed that the wife (I-2) was heterozygous for c.4006C>T, p.Gln1336*, for which the proband is homozygous. This being a non-complementary union, both their children (daughter II-1, son II-2) are also HI and are homozygous for the same variant (Fig. 3A).

This family reported postlingual progressive HL, a finding that is supported by their ability for verbal communication. The proband (I-1) and his daughter (II-1) have bilateral profound SNHL, while with hearing aids his son (II-2) (who died shortly after the study), had moderately-severe SNHL in the right ear and moderately-severe mixed HL in the left ear (Fig. 3D). Temporal lobe CT and MRI did not detect any gross abnormalities in the proband and his daughter; however, a slight hypo-pneumatization in the inferior part of the left mastoid bone air cells was observed in the son (II-2) (Fig. 3C). VNG results were normal in the daughter and son, while the proband had spontaneous nystagmus, saccade latency (rightward), delayed smooth pursuit and optokinetic nystagmus, which could be age related. vHIT indicated that all three HI subjects have low VOR in all the semicircular canals (SCCs) and hypoactive labyrinths, signifying vestibular dysfunction (Fig. 3E).

Family *DXN*CHM46 has autosomal dominant SNHL, with HI individuals in at least four generations (Supplementary Fig. 3). Exome sequencing of the proband (II-3) revealed multiple variants in known HL-associated genes. In order to identify the causative genotype, exome sequencing was therefore extended to six other family members, including four HI and two normal hearing individuals (Fig. 4A). This method identified a novel variant c.2962C>T, p.Pro988Ser in *SLC12A2* (MIM: 600840) (Fig. 4B). The gene has been recently associated with DFNA78 (MIM: 619081) in humans, in two studies (Morgan et al. 2020; Mutai et al. 2020). SLC12A2 is involved in sodium, potassium and chloride ion transport, and mouse model studies have shown that Slc12a2 plays a crucial role in endolymph homeostasis (Wangemann, 2006).

**Fig. 4.**
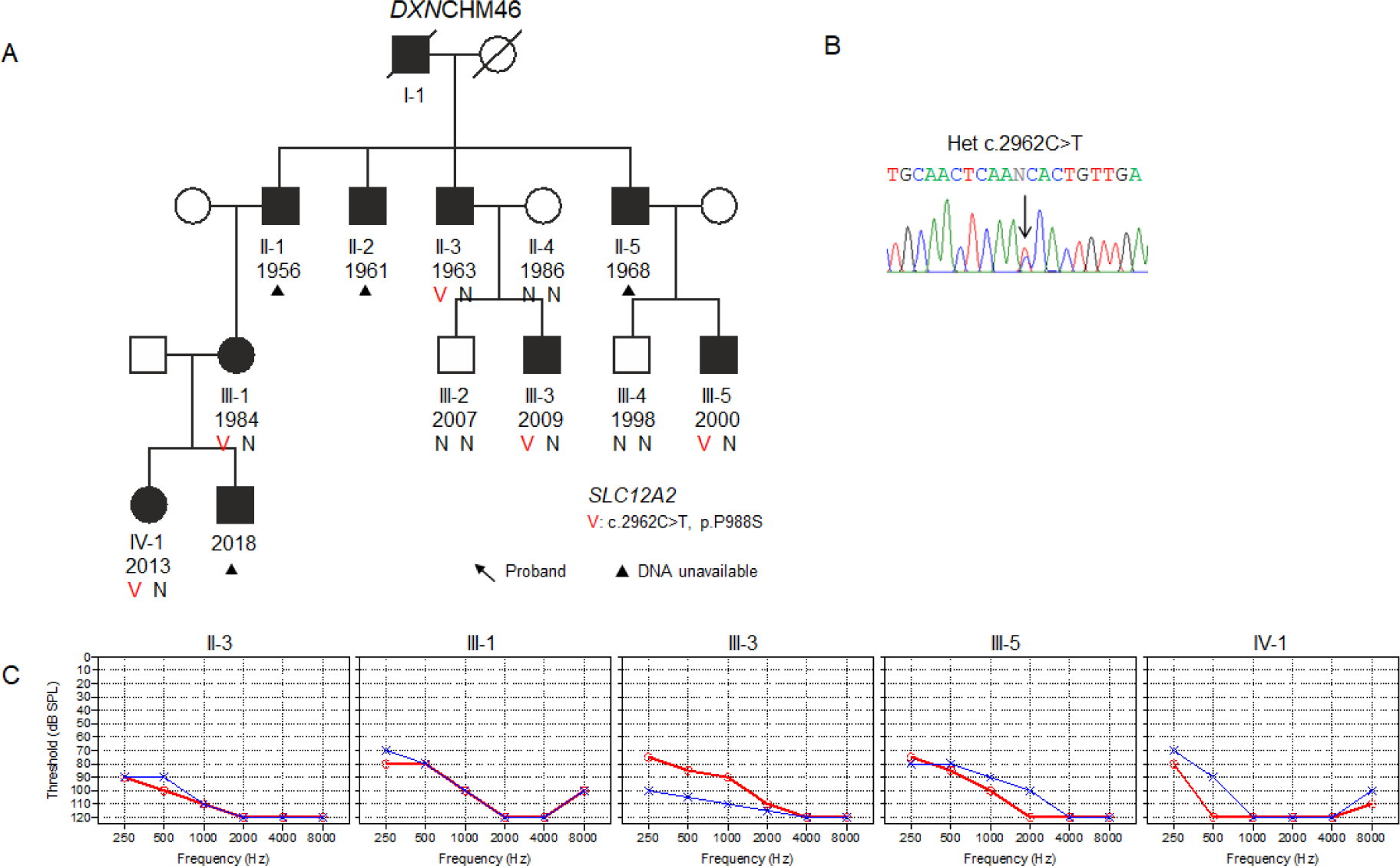
Segregation analysis and genotype-phenotype correlation in the *DXN*CHM46 family. The pedigree (A) shows the autosomal dominant HL segregating with the *SLC12A2* variant shown in the chromatogram (B). The audiograms (C) of II-3, III-1, III-3, III-5 and IV-1, show severe-to-profound HL

The novel variant c.2962C>T, p.Pro988Ser is located in exon 21 which encodes a part of the C-term intracellular domain of SLC12A2. Mutations in this exon have been reported to decrease the chloride influx activity of the protein (Mutai et al. 2020). p.Pro988Ser is predicted to decrease protein stability (Table 1), with a structural comparison of the <u>*wt*</u> and *mt* proteins predicting an RMSD of 1.81 and a TM score of 0.94.

This multiplex family has twelve HI individuals spanning across four generations, and segregating in an autosomal dominant mode (Supplementary Fig. 3). However, only five HI members of the family could be tested. The proband (II-3), who was found to have the *SLC12A2* heterozygous c.2962C>T, p.Pro988Ser variant has three HI brothers (II-1, II-2, II-5) who did not consent to participate. The proband’s unrelated, normal hearing wife (II-4) tested negative for the variant. This couple has two sons, one with normal hearing (III-2), who tested negative, and one with HI (III-3), who was heterozygous for c.2962C>T, p.Pro988Ser. Similarly, the proband’s HI nephew (III-5), HI niece (III-1) and niece’s HI daughter (IV-1), all harbour the heterozygous variant, while his normal hearing nephew (III-4) tested negative, clearly supporting the variant segregating with HL in this family (Fig. 4A).

Audiometry in the HI proband (II-3), his son (III-3), nephew (III-5), niece (III-1) and her daughter (IV-1) revealed prelingual bilateral severe-to-profound HL in these individuals (Fig. 4C). Furthermore, we were informed that the HI members of this family responded to sound until the age of 18 to 24 months, after which the hearing deteriorated.

## Discussion

Although numerous HL-associated genes have been identified in Indian families (Yan et al. 2015), the genetic landscape of HL in the population is not well understood and the unparalleled genetic and phenotypic heterogeneity has posed a challenge to the causal diagnosis of HL (Van Camp et al. 1997). In our previous study, four HI individuals were found to have DFNB12, with either homozygous or compound heterozygous *CDH23* variants (Vanniya et al. 2018).

In our present study, five unrelated HI probands from South Indian assortative mating families, with heterozygous *CDH23* variants, were subjected to exome sequencing to resolve the HL etiologies. Six variants were identified in four HL genes in these families. This is the first report to document HL-associated with *PNPT1*, *PTPRQ* and *SLC12A2* variants in the Indian population.

In family *DXD*CHE32, the proband (II-2) had compound heterozygous *PNPT1* variants c.137C>G, p.Ala46Gly/c.1619A>G, p.Asn540Ser, causing autosomal recessive, prelingual, bilateral SNHL (DFNB70) with vestibular dysfunction and unilateral progressive vision loss. No study has reported vestibular dysfunction in patients with DFNB70. *PNPT1* variants have been reported to cause multisystem dysfunction, including optic atrophy and auditory neuropathy (Alodaib et al. 2017). In one study, the co-occurrence of neurodegenerative features such as ataxia, dystonia and cognitive decline in the 4^th^ decade, further presenting with optic atrophy by the 6^th^ and 7^th^ decade, were reported (Eaton et al. 2018). Except for the unilateral progressive vision loss, none of the neurodegenerative features reported previously, were observed in the proband (who is in her 50s). In addition, the possibility of diabetes-associated vision loss could not be ruled out. Our results therefore support the proposition that *PNPT1* mutations cause a variable spectrum of phenotypic outcomes (Eaton et al. 2018).

The unrelated HI couple of the *DXD*PTM56 family, and their HI son, have DFNB3. While the husband (I-1) is homozygous for *MYO15A* c.5673C>G, p.Tyr1891* (novel), the wife harbors compound heterozygous *MYO15A* variants 4454T>C, p.Leu1485Pro/c.5673C>G, p.Tyr1891*, occurring in the myosin motor domain. The presence of the same novel variant in unrelated probands suggests that *MYO15A* variants may be relatively prevalent in South India. All the three HI tested had prelingual bilateral profound SNHL. While, DFNB3 is often associated with milder phenotypes, mutations in the myosin motor domain of myosin XVa have been strongly associated with severe phenotypes (Zhang et al. 2019). The genotype-phenotype correlation in this family substantiates the association of severe phenotype with myosin motor domain mutations.

In family *DXN*CHE29, a novel homozygous *PTPRQ* variant, c.4006C>T, p.Gln1336*, causing DFNB84A, was associated with postlingual progressive HL, in the proband (I-1) and his two HI children (II-1, II-2). However, although studies have associated DFNB84A with prelingual progressive HL (Schraders et al. 2010; Sakuma et al. 2015), the proband (I-1) and his two HI children (II-1, II-2) in the *DXN*CHE29 family reported postlingual progressive HL. The proband (I-1) and his daughter (II-1) have a severe phenotype of bilateral profound SNHL, while a milder phenotype was observed in his son (II-2), who used a hearing aid for his moderately severe SNHL in the right ear, and mixed HL in the left ear. A CT scan of the temporal lobe of the son (II-2) revealed a slight hypopneumatization of the left mastoid air cells, which agrees with previous results that conductive HL may be associated with poorly pneumatized mastoids (Sade 1992). These results suggest a need to evaluate the beneficial outcomes of hearing aids and the presence of conductive HL resulting from bone abnormalities in *PTPRQ*-associated HL.

The multiplex family *DXN*CHM46 has autosomal dominantly inherited prelingual HL (DFNA78), with a novel variant c.2962C>T, p.Pro988Ser in exon 21 of *SLC12A2* associated with severe-to-profound SNHL across three generations of the family. While the family members reported deterioration of hearing at around 2 years of age, suggesting a possibility of prelingual progressive HL, this could not be verified. Four *SLC12A2* variants, including three exon 21 variants, have been recently reported with DFNA78 in humans (Morgan et al. 2020; Mutai et al. 2020). Mutai et al. (2020) reported a variant (p.(Pro988Thr)) in the same residue as that observed in the *DXN*CHM46 family (p.Pro988Ser), suggesting that the residue Pro988 could be a mutational hotspot. Exon 21 in *SLC12A2* is critical for normal functioning of the protein, and mutations in the exon have been reported to exert a dominant negative effect (Mutai et al. 2020). The frequent occurrence of *SLC12A2* exon 21 variants in DFNA78 suggests that it could represent a region of mutational hotspots that should be a priority region for screening.

The present study illustrates the complexity of HL in these four assortative mating South Indian families. The allelic and genotypic heterogeneity observed in these families support the idea that complex genotypes and phenotypes are associated with HL. This study emphasizes the importance of clinical phenotyping, especially vestibular function tests, in order to provide genetic counselling with added management options.

Routine genetic diagnosis can be achieved with rapid, cost effective diagnostic strategies by prioritizing genes, hotspot screening or population-specific gene panel tests. However, there remains a need for large cohort studies in order to better understand the ethnicity-specific genetic landscapes that will eventually lead to effective genetic diagnosis in HL.

## Data Availability Statement

The novel variants identified in the study have been submitted to ClinVar (https://www.ncbi.nlm.nih.gov/clinvar/), and the accession numbers are: SCV001366119 (https://www.ncbi.nlm.nih.gov/clinvar/variation/973493/), SCV001366122 (https://www.ncbi.nlm.nih.gov/clinvar/variation/973491/), SCV001366123 (https://www.ncbi.nlm.nih.gov/clinvar/variation/972899/).

## Authors’ contributions

CRS conceived the presented idea and supervised the project. CRS and PVS planned the experiments. JC, PVS, and JMJ recruited the subjects and collected samples for this study in the field. KBA, NS, and LC facilitated the exome sequencing and data analysis. TR and ZB performed the exome data analysis. PVS performed the molecular biology experiments and *in silico* analysis, collected clinical and molecular data, and interpreted the results. MK validated the genotype of a proband in the *DXD*PTM56 family. PVS and CRS drafted the manuscript, with inputs by KBA. All the authors read, reviewed and approved final version of the manuscript.

## Acknowledgements

This study was supported by research grants sanctioned to CRS from University Grants Commission, India (F. No.37-443/2009(SR)); Ad-Hoc Research Project (5/8/10-17(Oto)/CFP/2011-NCD-I) from the Indian Council of Medical Research, India; Department of Biotechnology, Multicentric Project, India (BT/PR26850/MED/12/80/2017); Indian Council of Medical Research- Senior Research Fellowship, India (No.45/1/2016-HUM-BMS); Department of Science and Technology- Women Scientist Scheme-A (SR/WIS-A/LS-390/2018(G)); University Grants Commission- Special Assistance Programme; Department of Science and Technology- Fund for Improvement of S&T Infrastructure (II); and University Grants Commission- Universities with Potential for Excellence (Phase II).

We are grateful to all the participants for their consent and co-operation. We thankfully acknowledge Dr. A. Pavithra for her participation in the field work and sharing with us the DFNB1 status for all the study subjects. We thank Dr. Mohan Kameswaran and Dr. Sudha Maheshwari, Madras ENT Research Foundation, Chennai for facilitating audiometry, VNG and VHIT in the study subjects. We thank Dr. R. Rajeswaran, Sri Ramachandra Medical Centre, Chennai, for the CT and MRI interpretations.

---

**Supplementary Fig. 1** PCR-RFLP analysis for screening the variants in controls

**Supplementary Fig. 2** Pedigree of family *DXN*CHE29. Black-filled symbols represent individuals with HL; grey-filled symbols represent individuals with known postlingual progressive HL

**Supplementary Fig. 3** Pedigree of family *DXN*CHM46. Black-filled symbols represent individuals with HL

