## Supplemental files for "Genetic and phenotypic heterogeneity in *PNPT1*, *MYO15A*, *PTPRQ* and *SLC12A2* variants detected among hearing impaired assortative mating families in Southern India"

**Running Title**

### Genetic and phenotypic heterogeneity of hearing impaired in Southern India

Paridhy Vanniya. S[1], Jayasankaran Chandru[1], Justin Margret Jeffrey[1], Tom Rabinowitz[2], Zippora Brownstein[3], Mathuravalli Krishnamoorthy[1], Karen B. Avraham[3], Le Cheng[4], Noam Shomron[2], C. R. Srikumari Srisailapathy[1]*

[1]Department of Genetics, Dr. ALM PG Institute of Basic Medical Science, University of Madras, Chennai- 600 113, India

[2]Department of Cell and Developmental Biology, Sackler Faculty of Medicine, Tel Aviv University, Tel Aviv- 6997801, Israel

[3]Department of Human Molecular Genetics and Biochemistry, Sackler Faculty of Medicine and Sagol School of Neuroscience, Tel Aviv University, Tel Aviv- 6997801, Israel

[4]BGI Genomics, Shenzhen- 518083, China

**Correspondence**

*C. R. Srikumari Srisailapathy, Department of Genetics, Dr. ALM PG Institute of Basic Medical Science, University of Madras, Chennai-600 113, India. Telephone: 91-44-24547066;

**Supplementary Table 1** Primers used for confirmation and screening the variants by Sanger sequencing

| Primers | Sequence (5' – 3') | Tm (ºC) | Amplicon (bp) |
|---|---|---|---|
| *PNPT1* exon 1, Forward | CCAGGAGGTGTTGACACAGA | 62 | 494 |
| *PNPT1* exon 1, Reverse | CAGAGGCTAAAGCCCAACAG | 62 | |
| *PNPT1* exon 20, Forward | CTCAGGCATTTATGAGGGAAA | 61 | 363 |
| *PNPT1* exon 20, Reverse | TGAAGGGAGAATCAAGCACA | 61 | |
| *MYO15A* exon 11, Forward | AGGCCACCACACTACTGGTC | 62 | 333 |
| *MYO15A* exon 11, Reverse | GAAACAGAGAAGGCCTTGGA | 64 | |
| *MYO15A* exon 23, Forward | CCTATTCTGTCTCCACGGACTT | 64 | 525 |
| *MYO15A* exon 23 Reverse | TCTCAGGTCCCTGAAATGC | 60 | |
| *PTPRQ* exon 24, Forward | AGCCAAACTTGTTGGACTGG | 60 | 339 |
| *PTPRQ* exon 24, Reverse | AGCCAAACTTGTTGGACTGG | 62 | |
| *SLC12A2* exon 2,1 Forward | GCCACAGTTCAACATCTTCTCA | 60 | 621 |
| *SLC12A2* exon 21, Reverse | CCGGCTGTCTGGGTCTAATA | 60 | |

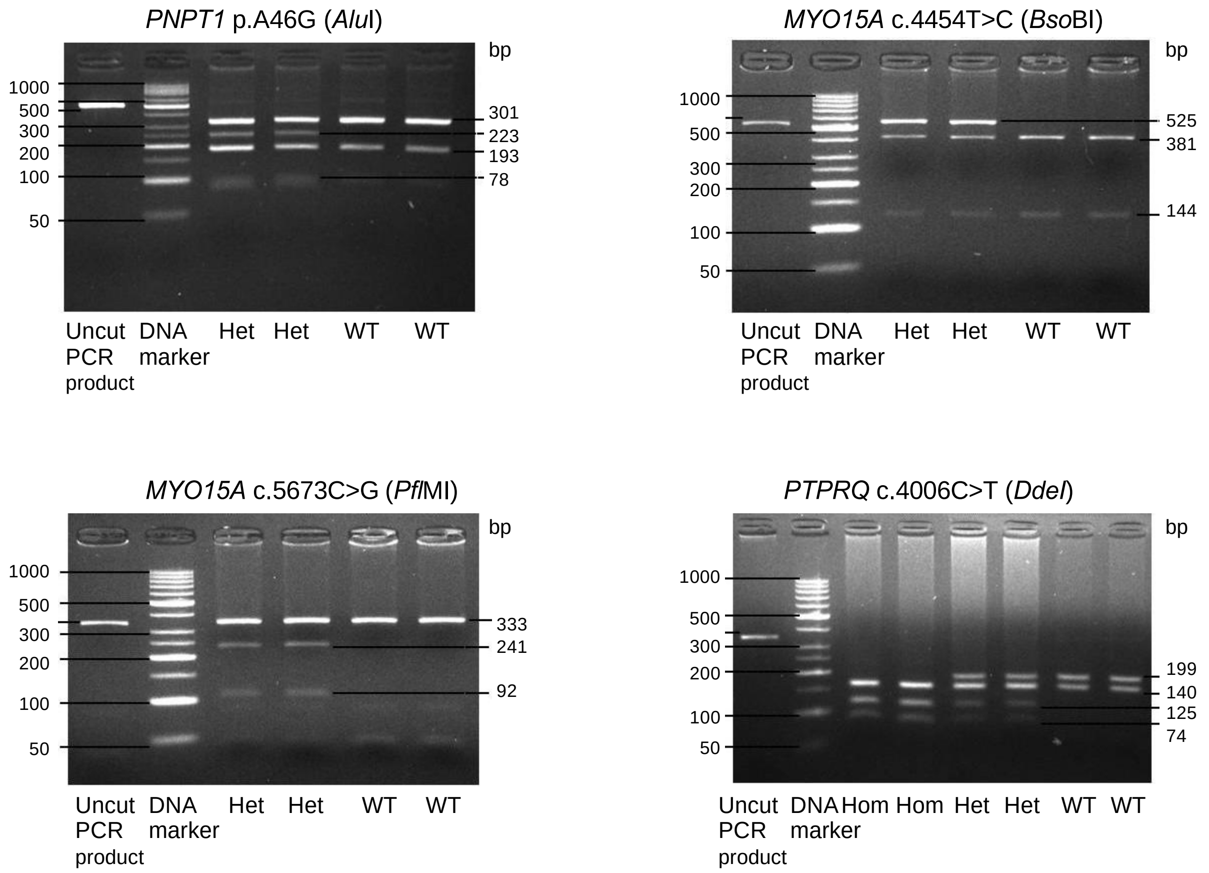


**Supplementary Fig. 1** PCR-RFLP analysis for screening the variants in controls.

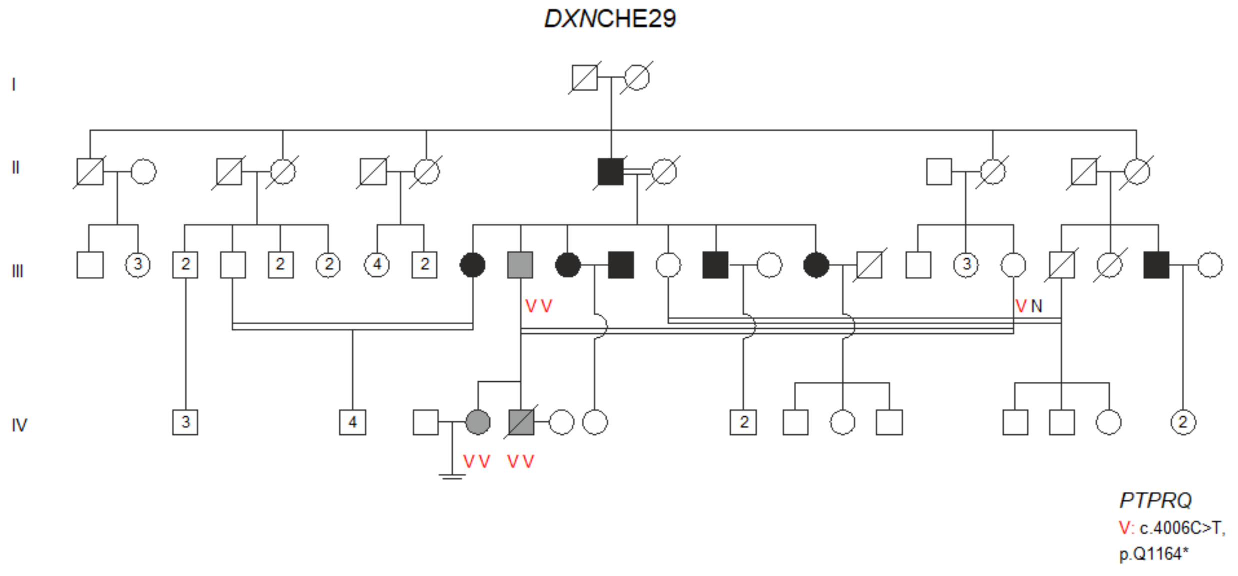


**Supplementary Fig. 2** Pedigree of family *DXN*CHE29. Black-filled symbols represent individuals with HL; grey-filled symbols represent individuals with postlingual progressive HL

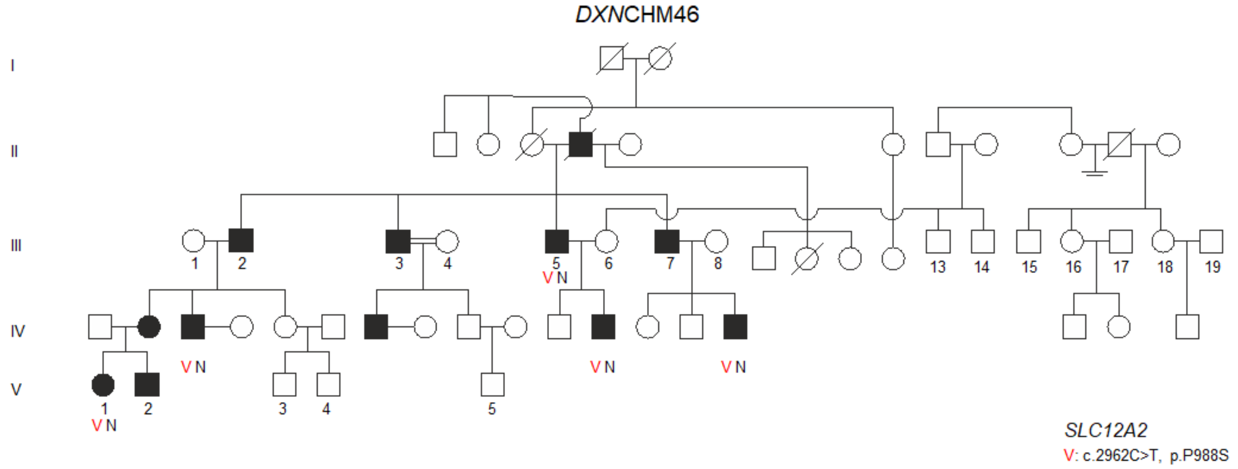


**Supplementary Fig. 3** Pedigree of family *DXN*CHM46. Black-filled symbols represent individuals with HL
